# The discovery of phages in the Substantia Nigra and its implication for Parkinson’s Disease

**DOI:** 10.1101/2024.07.13.603353

**Authors:** Yun Zhao, Changxian Xiong, Bingwei Wang, Daotong Li, Jiarui Liu, Shizhang Wei, Yujia Hou, Yuan Zhou, Ruimao Zheng

## Abstract

**Background:** A century ago, a mystery between virus and Parkinson’s disease (PD) was described. Owing to the limitation of human brain biopsy and the challenge of electron microscopy in observing virions in human brain tissue, it has been difficult to study the viral etiology of PD. Recent discovery of virobiota reveals that viruses coexist with humans as symbionts. Newly-developed transcriptomic sequencing and novel bioinformatic approaches for mining the encrypted virome in human transcriptome make it possible to study the relationship between symbiotic viruses and PD. Nevertheless, whether viruses exist in the human substantial nigra (SN), and whether symbiotic viruses underlie PD pathogenesis remain unknown.

**Methods:** We collected current worldwide human SN transcriptomic datasets from the United States, the United Kingdom, the Netherlands and Switzerland. We used bioinformatic approaches including viruSITE and the Virus-Track to identify the existence of viruses in the SN of patients. The comprehensive RNA sequencing-based virome analysis pipeline was used to characterize the virobiota in the SN. The Pearson’s correlation analysis was used to examine the association between the viral RNA fragment counts (VRFC) and PD-related human gene sequencing reads in the SN. The differentially expressed genes (DEGs) in the SN between PD patients and non-PD individuals were used to examine the molecular signatures of PD and also evaluate the impact of symbiotic viruses on the SN.

**Findings:** We observed the existence of viruses in the human SN. A dysbiosis of virobiota was found in the SN of PD patients. A significant correlation between VRFC and PD-related human gene expression was detected in the SN of PD patients. These PD-related human genes correlated to VRFC were named as the virus-correlated PD-related genes (VPGs). We identified three bacteriophages (phages), including the Proteus phage VB_PmiS-Isfahan, the Escherichia phage phiX174 and the Lactobacillus phage Sha1, that might impair the gene expression of neural cells in the SN of PD patients. The Proteus phage VB_PmiS-Isfahan was a common virus in the SN of patients from the UK, the Netherlands, and Switzerland. VPGs and DEGs together highlighted that the phages might dampen dopamine biosynthesis and weaken cGAS-STING function.

**Interpretation:** This is the first study to discover the involvement of phages in PD pathogenesis. A life-long low symbiotic viral load in the SN may be a contributor to PD pathogenesis. Our findings unlocked the black box between brain virobiota and PD, providing a novel insight into PD etiology from the perspective of phages-human symbiosis.

## Introduction

Parkinson’s disease (PD) is characterized by loss of dopaminergic neurons in the substantia nigra (SN), involuntary shaking and muscle rigidity^1–3^. Pathological hallmarks of PD include neuroinflammation^4–7^, imbalanced protein homeostasis^8–11^, oxidative stress^12–15^, cell aging^16^, regulation of neurotransmitter and neuronal apoptosis^17,18^. An estimated 90% of PD patients are known to be caused by environmental factors^19–21^. Since the 1918 influenza pandemic, association between viruses and PD has been debated^22–28^. For a century, clinical evidence reveals that virus may cause PD-like symptoms^29,30^. Incidence of PD was lower in patients who received antiviral therapy^31,32^. Thus, there is a need to study the mechanism by which virus may affect PD pathogenesis.

In recent years, discovery of virobiota reveals that viruses coexist with humans as symbionts^33–42^. Human body is colonized with substantial communities of viruses^43^, termed ‘virobiota’^44^. Bacteriophages (phages) are the most abundant viral entities in humans and also the major component of intestinal virobiota^45–47^. Phages can be disseminated throughout human tissues^48^, and can cross blood-brain barrier (BBB) to access brain^49–54^. In mammalian cells, phages can enter organelles, maintain long-term residence and affect cell function^55–58^. Phage genomes can be integrated into human chromosomes^59^, and phage gene transcripts can be detected in human cells^60,61^. In the brain, phages can enter neural cells to cause neuroinflammation or neuronal death^55,56,62,63^. Together, these studies suggest that viruses/phages may affect function of mammalian cells. However, whether viruses/phages exist in the human SN; and whether viruses/phages may be linked to PD pathogenesis remain unknown.

Next-generation sequencing becomes novel approach to identify and characterize virobiota in tissues^64^. Sequencing of viral mRNA fragments that encrypted in human transcriptome reveal viral involvement in human diseases^65^. Newly-developed bioinformatic approaches, viruSITE^66^ and Virus-Track^64^ are designed for mining the encrypted virome in transcriptome of host tissues. Correlation analysis between viral RNA fragment counts (VRFCs) and human host gene sequencing reads can uncover the relationship between virus and human diseases^67,68^. Genome-wide microarray datasets of the human SN also contributes to characterize PD-related gene expression^69–74^. Therefore, worldwide RNA-sequencing/microarray studies performed on autopsy SN samples of PD patients can provide resources for exploring the link between the SN virobiota and PD pathogenesis. In this study, by using transcriptomic datasets of SN samples from Geneva University Hospitals, Netherlands Brain Bank, and Parkinson’s UK Brain Bank, we identified a viral existence in the SN, and a strong correlation between VRFCs of phages and PD-related human gene expression in the SN of PD patients. Our findings discovered that brain virobiota may underlie PD pathogenesis.

## Methods

### Prevention of microbial RNA and RNase contamination

To prevent microbial RNA or RNA enzyme contamination, sterile procedures for the SN dissection and collection were performed. For detailed information on the procedures of the SN dissection and collection, please refer to the brain bank websites: Netherlands Brain Bank (https://www.brainbank.nl/brain-tissue/autopsy/); the Parkinson’s UK Brain Bank (https://www.parkinsons.org.uk/research/parkinsons-uk-brain-bank), and the Geneva University Hospitals (https://www.hug.ch/en/clinical-pathology).

### Flow chart of bioinformatic analysis pipelines was shown in Fig 2. Data Collection

We used “Parkinson’s disease” as keywords to search for genome-wide expression studies in NCBI-GEO (http://www.ncbi.nlm.nih.gov/geo/) and European Genome-phenome Archive (EGA) platform (https://ega-archive.org/).The inclusion criteria included as follows:

(1) The studies that were designed for exploring gene expression in the SN for PD patients and non-PD individuals were the first choice for inclusion.
(2) The study type was Gene Expression Profiling by RNA-seq or microarray.
(3) The microarray studies which comprised of cell intensity file (CEL) raw files. Besides, to reduce the bias from different microarray platform, only data from two widely used platforms Affymetrix Human Genome U133A and Affymetrix Human Genome U133 Plus 2.0 were considered.

The raw RNA-seq data were retrieved from NCBI SRA database (https://www.ncbi.nlm.nih.gov/sra) or generously shared by Prof. Paul Lingor and Dr. Lucas Caldi Gomes on EGA platform.

We then performed analysis of five RNA-seq datasets (EGAD00001006883, GSE169755, GSE114918, GSE136666, and GSE114517) and five microarray datasets (GSE20141, GSE49036, GSE7621, GSE8397 and GSE20292). Details of the datasets are provided in eTable 1. The information of PD patients and non-PD individuals in this study are provided in eTable 2.

### Analysis of virome composition and structure in the SN

To identify viral fragments from the RNA-seq data of the SN, the raw FASTQ sequencing reads were first pre-processed by fastp software (version 0.21.0, default parameters)^75^ for quality control and adaptor trimming. Then the reads were aligned to a merged reference genome file combining human reference genome (hg38) and a comprehensive collection of 13559 virus genomes from viruSITE database (http://www.virusite.org/index.php, version 2021.2)^66^ by using the STAR software (version 2.7.8)^76^. When running the STAR software, we adopted the parameters recommended by Viral-Track^64^, which is a recently established computational pipeline for detecting viral reads from sequencing data. Based on the reads aligned to viral genome, viral read counts were obtained via Viral-Track to assess the abundance of each virus, i.e. VRFC. Instead of the built-in thresholds of Viral-Track which were designed for the near full-length viral gene detection purpose^64^, two alternative criteria were applied for false positive control for our purpose of viral fragment detection:

(1) All viral reads should assemble short viral contigs. For each sample, all reads aligned to a virus genome were extracted by SAMtools (version 1.12)^77^ and submitted to the Trinity contig assembly pipeline (version 2.13.2)^78^ using the virus genome as the contig assembly reference and allowing no intron inside the contigs. Only assembled contigs longer than 50 nucleotides and showing at least 90% identity to the reference genome was retained.
(2) All viral reads considered in virus quantification should NOT be aligned to any chromosome of the human refence genome.

To further assess the composition and structure of the SN virome, the viral read counts or the normalized viral RPKM (reads per kilobase per million reads mapped) were imported into R (version 4.0.2), as per the requirement of the software used in the subsequent analysis. Virus abundance, alpha diversity and beta diversity were calculated with R package phyloseq (version 1.32.0) and vegan (version 2.6.4). Permutational multivariate analysis of variance (PERMANOVA) implemented in vegan package was performed for Bray-Curtis dissimilarity. Data visualization was performed by R packages ggplot2 (version 3.4.0) and aPCoA (version 1.3). The statistically significant differences between PD and non-PD groups were determined by two-tailed Wilcoxon test using ggpubr (version 0.4.0) and ggsignif (version 0.6.3) R packages, and a *p* value < 0.05 was considered statistically significant.

### Human gene expression quantification and its correlation with viral gene expression

Raw FASTQ reads were preprocessed and aligned to human genome using the same method as above. Then, based on the reads aligned to the known genes in human reference genome, the human gene expressions were quantified by the featureCounts method of Rsubread R package (version 2.4.3)^79^, using the standard Ensembl gene annotation reference (http://www.ensembl.org/, version 104) and default parameters. Pearson’s correlation between virus expression and human gene expression in PD and non-PD patients were calculated respectively using the cor.test function in R. We focused on the correlations regarding to PD-related pathological genes. The PD-related human genes correlated to VRFC were termed VPGs. The correlation heatmap was generated using the pheatmap R package (version 1.0.12). The paired box plot was generated by the ggpubr R package (version 0.4.0) and the Wilcoxon signed-rank test was implied by the ggsignif R package (version 0.6.4).

### Differential expression analysis and functional enrichment analysis

Both RNA-seq and microarray-based gene expression profiles were considered in the differential expression analysis. To be scalable to the microarray data, the gene expression values from RNA-seq data were firstly transformed to log_2_(x+1). As for the microarray data, the raw CEL files were processed using the Robust Multichip Average (RMA) method for background correction and normalization, which is implemented in the affy (version 1.68.0) and gcrma (version 2.62.0) R packages. After removing duplicated gene probes and unspecific probes, all probes were mapping to single Entrez Gene IDs according to the corresponding probe annotation files. Data from RNA-seq and Microarray datasets were merged based on their shared genes, resulting a gene expression matrix covering 12,180 genes. Batch effects were supervised by principal component analysis (PCA) method and removed using the ComBat function of the sva R package (version 3.38.0). Negative expression values introduced during batch effect removal were truncated to zero. Differential gene expression analysis was carried out by the limma R package (version 3.46.0), and genes with a *p* value < 0.05 and |log2 (Fold change) | > 1 were considered significant differentially expressed genes (DEGs). Gene Ontology (GO) and Kyoto Encyclopedia of Genes and Genomes (KEGG) analysis for these DEGs were based on the Metascape database (http://metascape.org/gp/index.html#/main/step1), and a functional term with corrected *p* value < 0.05 was considered significantly enriched. The expression heatmap of all DEGs was plotted using pheatmap R package (version 1.0.12). GO biological processes network of DEGs was carried out by BiNGO^80^ tool in Cytoscape software^81^.

### Protein-protein interaction (PPI) network construction and comparison of the top hub interacting node genes

The STRING 11.5 database was used to predict the interactions of DEGs and virus-related genes identified in the study and to map the protein-protein interaction (PPI) network^82^. We selected the data analysis mode and default PPI confidence threshold of the STRING database to construct the PPI network. The protein networks were visualized by Cytoscape software^81^, and analyzed by the Network Analyzer tool based on degree. The degree indicates the number of interactions of each protein. We compared the relative expression of the DEGs across five RNA-seq datasets in this study. To assess the diagnostic value of DEGs, we compared the expression of the hub node genes (i.e., genes with interactions > 5). A receiver operating characteristic (ROC) curve was performed, and the ROC curve

## Results

### Characteristics of RNA-sequencing/microarray datasets of the SN of PD patients

Worldwide PD brain RNA-sequencing datasets from the United Kingdom (UK), the Netherlands, and Switzerland were collected (Figure 1); microarray-datasets from the United States (US), the UK and the Netherlands were enrolled (eTable 2). There are comparable PD prevalence, average life expectancy and gender composition among these countries. To ensure comparability and homogeneity of raw data, ‘Combat’ method was used to correct batch effects (Figure 2).

**Figure 1:**
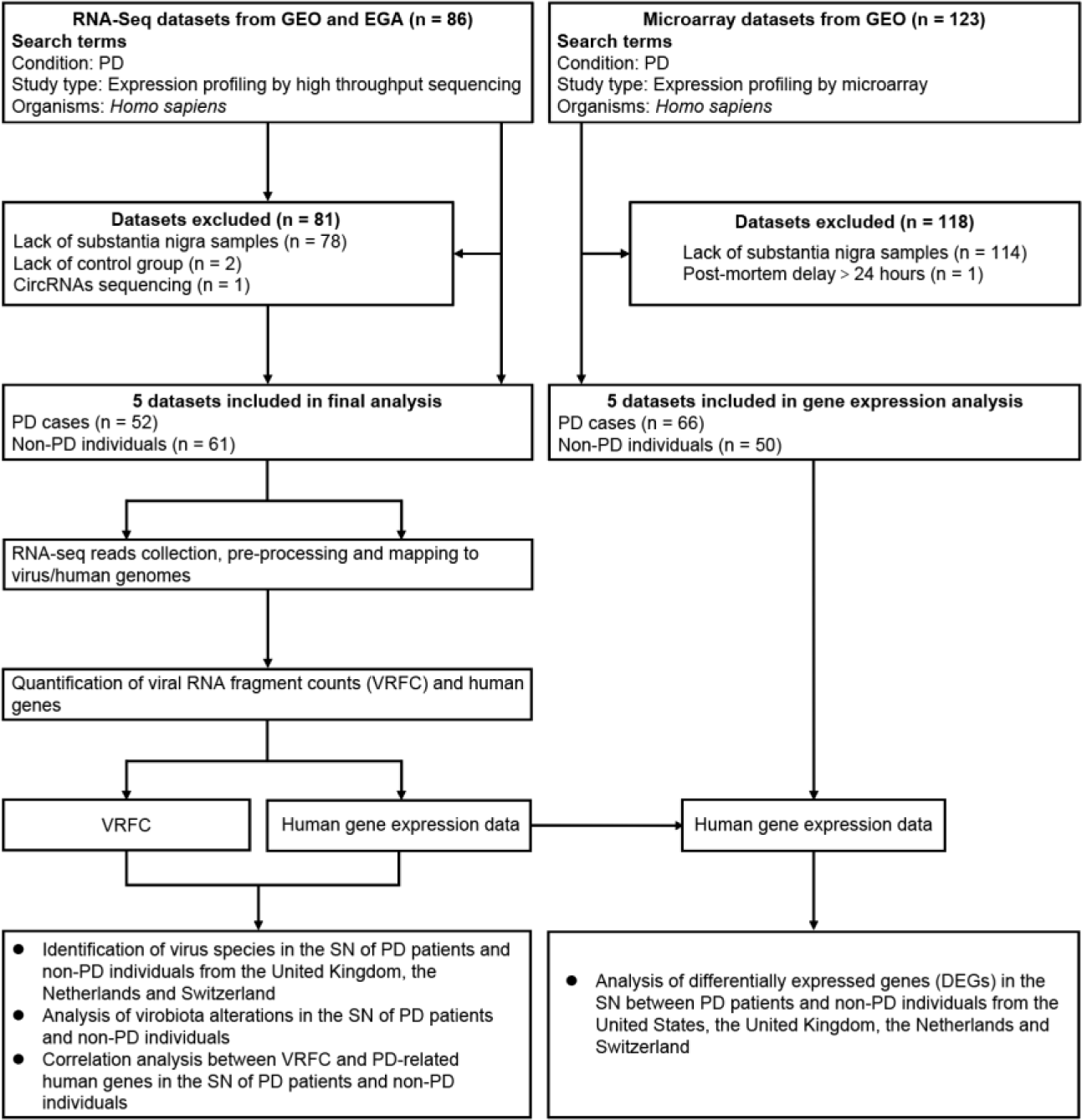
Enrollment and analysis of datasets.

**Figure 2:**
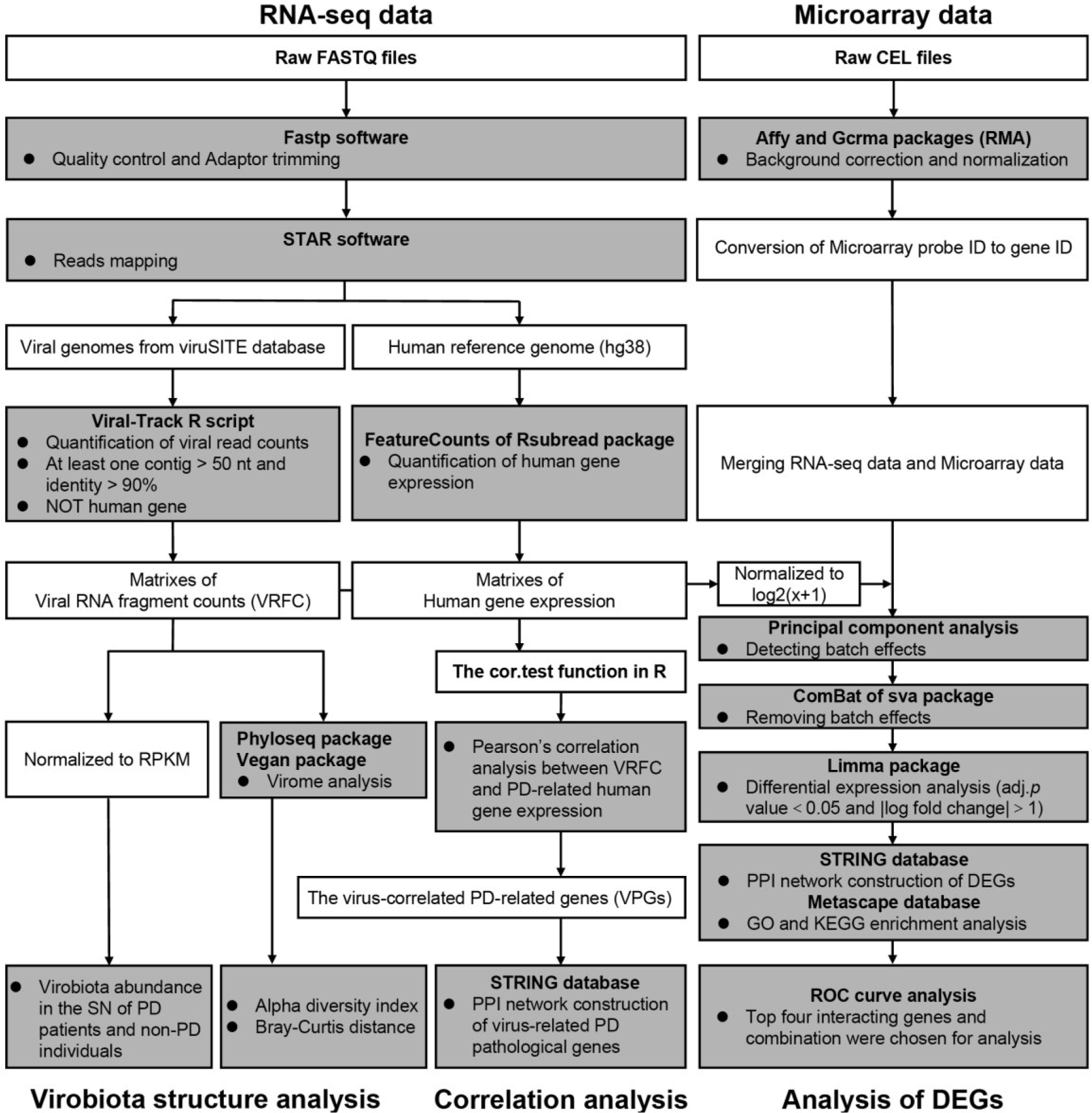
Flow chart of bioinformatic analysis pipelines.

### Viral sequences were detected in the human SN

Viral mRNA fragments were detected in the SN samples. These viruses included the phages that host gut microbiota, the viruses that host primates, and the viruses that host plants and arthropods (Table 1 and eTable 3). Proteus phage VB_PmiS-ISfahan and Escherichia phage Lambda ev017 were common viruses in the SN of patients from these countries, showing the viral population distribution may be geographically related (Figure 3). These findings revealed an existence of viruses in the human SN, suggesting that the human SN may be colonized by commensal virobiota.

**Figure 3:**
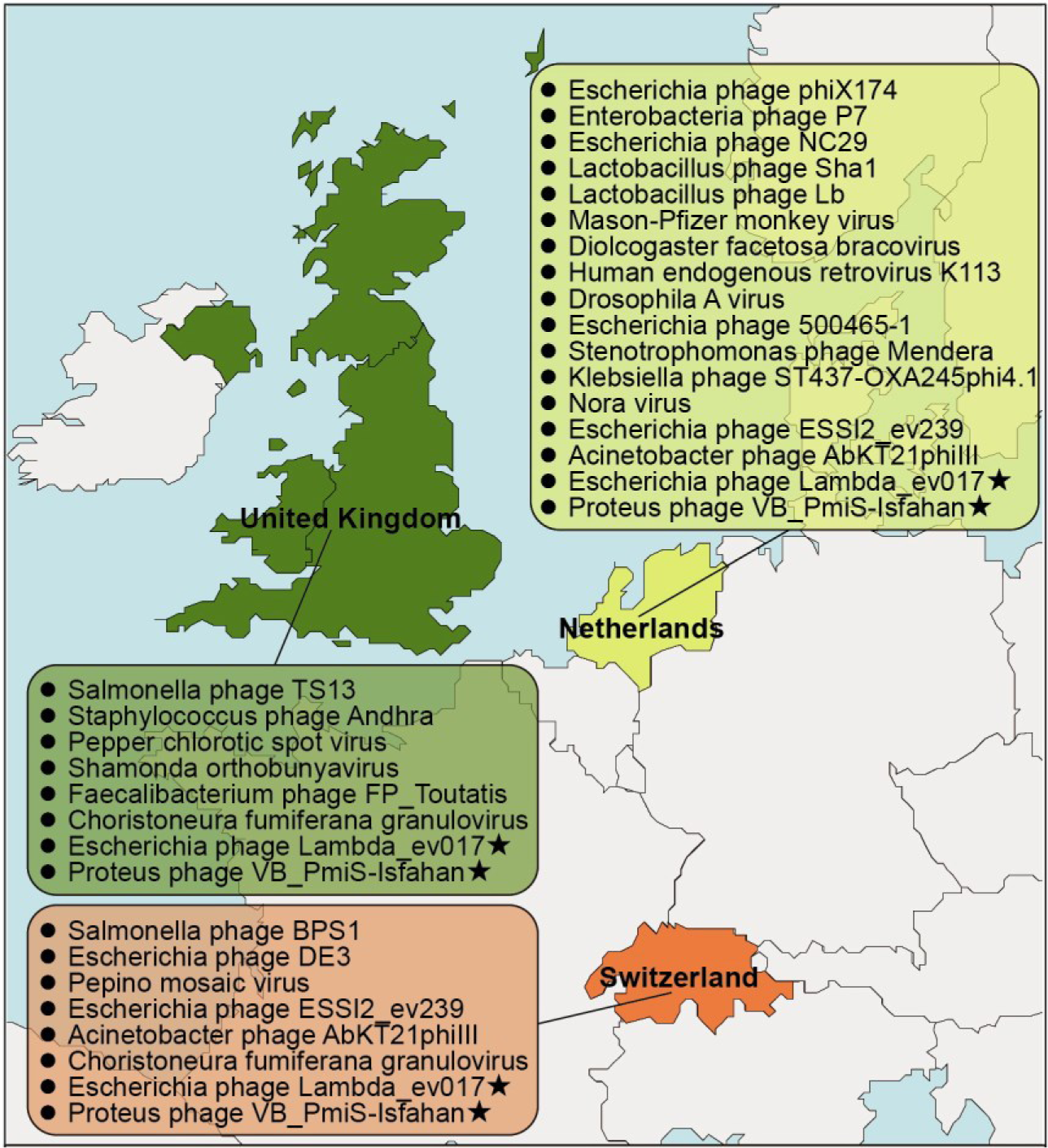
Identification of viral species in the SN samples from the brain banks of the UK, the Netherlands, and Switzerland.

**Table 1:** Viruses of nearest match identified in th SN of PD patients.

| Table 1 Viruses of nearest match identified in the substantia nigra of PD patients. |  |  |  |  |  |
| --- | --- | --- | --- | --- | --- |
| Viral family | Virus of nearest match | % Nucleotide identity | Contig size range (nt) | Main reported host | Baltimore classification |
| Siphoviridae | Escherichia phage Lambda_ev017 | 100 | 35-79 | Escherichia | I: dsDNA |
| Siphoviridae | Proteus phage VB_PmiS-Isfahan | 100 | 50-510 | Proteus mirabilis | I: dsDNA |
| Siphoviridae | Lactobacillus phage Sha1 | 100 | 53-122 | Lactobacillus sp. | I: dsDNA |
| Siphoviridae | Escherichia phage DE3 | 100 | 52-150 | Escherichia coli BL21(DE3) | I: dsDNA |
| Casjensviridae | Salmonella phage TS13 | 100 | 43-68 | Salmonella enterica | I: dsDNA |
| Casjensviridae | Salmonella phage BPS1 | 97 | 33-52 | Salmonella enterica subsp | I: dsDNA |
| Myoviridae | Faecalibacterium phage FP_Toutatis | 100 | 47-117 | Faecalibacterium prausnitzii | I: dsDNA |
| Peduviridae | Escherichia phage ESS12_ev239 | 100 | 35-93 | Escherichia | I: dsDNA |
| Peduviridae | Escherichia phage 500465-1 | 100 | 87-364 | Escherichia coli | I: dsDNA |
| Peduviridae | Klebsiella phage ST437-OXA245phi4.1 | 100 | 45-134 | Klebsiella pneumoniae | I: dsDNA |
| Retroviridae | Human endogenous retrovirus K113 | 100 | 53-136 | Homo sapiens | VI: ssRNA-RT |
| Retroviridae | Mason-Pfizer monkey virus | 100 | 61-68 | Macaca mulatta | VI: ssRNA-RT |
| Microviridae | Escherichia phage phiX174 | 100 | 53-5386 | Escherichia coli | II: ssDNA (+) |
| Microviridae | Escherichia phage NC29 | 99 | 92 | Escherichia | II: ssDNA (+) |
| Autographiviridae | Acinetobacter phage AbKT21phill | 94 | 52-63 | Acinetobacter baumannii | I: dsDNA |
| Rountreeviridae | Staphylococcus phage Andhra | 95 | 29-56 | Acanthamoeba castellanii | I: dsDNA |
| Polydnaviriformidae | Diolcogaster facetosa bracovirus | 100 | 55-93 | Glyptapanteles flavicoxis | I: dsDNA |
| Baculoviridae | Choristoneura fumiferana granulovirus | 100 | 36-77 | Choristoneura fumiferana | I: dsDNA |
| Tospoviridae | Pepper chlorotic spot virus | 93 | 71 | Peppers | V: ssRNA (-) |
| Peribunyaviridae | Shamonda orthobunyavirus | 94 | 87 | Bos taurus | V: ssRNA (-) |
| Alphaflexiviridae | Pepino mosaic virus | 96 | 67 | Solanum lycopersicum | IV: ssRNA (+) |
| NA | Enterobacteria phage P7 | 100 | 35-360 | Escherichia coli | I: dsDNA |
| NA | Stenotrophomonas phage Mendera | 96 | 56 | Stenotrophomonas maltophilia | I: dsDNA |
| NA | Lactobacillus phage Lb | 100 | 73-116 | Lactobacillus brevis ATCC 367 | I: dsDNA |
| NA | Nora virus | 98 | 84-113 | Drosophila melanogaster | IV: ssRNA (+) |
| NA | Drosophila A virus | 99 | 77-176 | Drosophila melanogaster | IV: ssRNA (+) |

### Dysbiosis of virobiota in the SN of PD patients

In SN samples, 11 viral families were detected (Figure 4A). In PD patients, viral families Peduoviridae, Microviridae and Autographiviridae were enriched, whereas viral family Siphoviridae was diminished, as compared with non-PD individuals (Figure 4B). To explore difference in composition of the SN virobiota between groups, we quantified the presence ratio of core, common and unique viral species (corresponding to viral species shared among > 80%, 30 -80% and < 30% of the individuals, respectively). In the SN of PD patients, the core species accounted for higher proportion, whereas unique species was diminished (Figure 4C). Richness (Chao1) and diversity (Shannon) of virome in the SN did not differ between groups (Figure 4D and E). Composition of virome in the SN of two groups was separated into two distinct clusters. Viral community dissimilarity among PD patients was higher than that of non-PD individuals (Figure 4F and G). Analysis of differentially present taxa at the species level show a remarkable difference in viral community structures between groups (Figure 4H). Together, these findings uncovered a dysbiosis in the SN virome of PD patients, suggesting that altered core species proportion and virobiota dysbiosis may be associated with PD pathogenesis.

**Figure 4:**
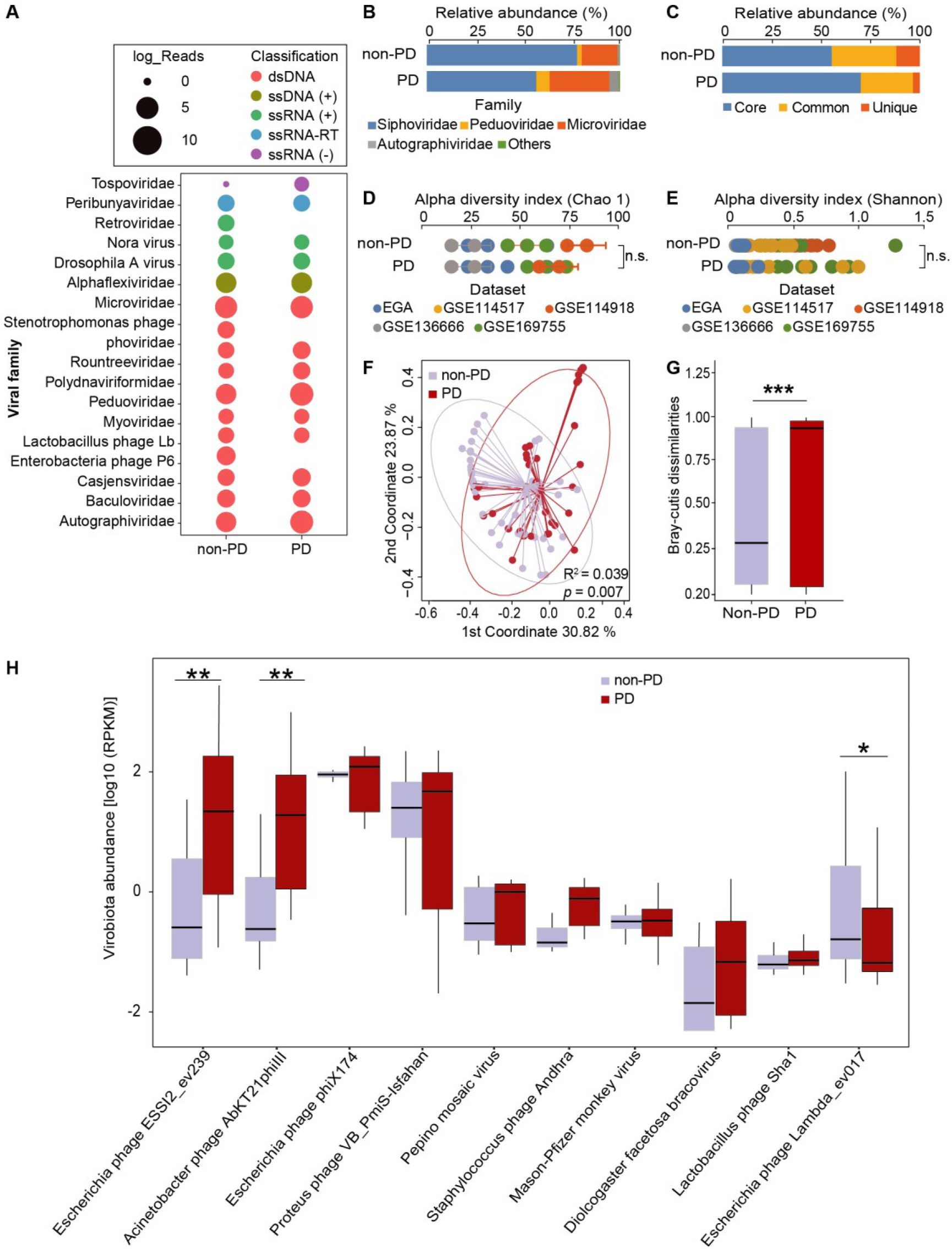
Dysbiosis of virobiota in the SN of PD patients. (A) Bubble plot showing the read abundances of different viral families in the SN virome. Read counts are log2-transformed and represented by the circle. (B) The proportion of viral families in the SN virome. (C) The proportion of core, common and unique viral species in the SN virome. The core species, common species and unique species correspond to viral species shared among > 80%, 30% – 80% and < 30% of studied group, respectively. (D to E) Comparison of α-diversity of the SN virome based on Chao1 richness index (D) and Shannon diversity (E). n.s., not significant. Statistical significance was determined by t-test. (F) Principal Coordinate Analysis (PCoA) plot of the Bray-Curtis distance showing the PD patients and non-PD individuals. Statistical significance for the Bray-Curtis distance was determined by PERMANOVA with permutations done 999 times. (G) Comparison of within-group SN virome Bray-Curtis dissimilarities between PD patients and non-PD individuals. Statistical significance was determined by t-test. \*\*\**p* value < 0.001. (H) Differential viral taxa between PD and non-PD at the species level. Differentially enriched viral species were determined by DESeq analysis. For viral abundance box plots, the boxes extend from the first to the third quartile (25th to 75th percentiles), with the median depicted by a horizontal line. RPKM, reads per kilobase per million mapped reads. \**p* value < 0.05 and \*\**p* value < 0.01.

### A strong negative correlation between VRFC of Proteus phage VB_PmiS-Isfahan and human PD-related gene sequencing reads in the SN of PD patients

To explore whether gene activity of symbiotic viruses in the SN may underlie PD pathogenesis, correlation analysis between VRFCs and PD-related human gene expression was performed (Figure 5). Based on GSE114517 dataset, a strong negative correlation between VRFCs of phages and PD-related human gene expressions (tyrosine hydroxylase, TH, a dopamine synthetase, etc.) was detected in the SN of PD patients; pathways-affected by these phages were similar in PD pathophysiology, although the phages belong to different genera and families (Supplementary Figure 2A, B and C. eTable 4, 5 and 6). We named these human PD-related genes which are correlated to the VRFC as “the virus-correlated PD-related genes (VPGs)”. The VPGs suppressed by the phages were enriched for the PD-related pathways including cGAS-STING response, oxidative stress, apoptosis, etc. (Supplementary Figure 2D and E. eTable 7). PPI network revealed 322 pairs of interactions among these VPGs (Supplementary Figure 1A). Top 19 VPGs with more than 15 interactions were shown in bar plot (Supplementary Figure 1B). A strong correlation coefficient among the VPGs was observed in PD patients (Supplementary Figure 1C). Venn analysis revealed a large overlap of 1313 genes among the VPGs affected by three phages (Supplementary Figure 2F). Proteus phage VB_PmiS-Isfahan was a common virus in the SN of patients from these countries. Likewise, based on all 5 datasets, similar results between VRFCs of Proteus phage VB_PmiS-Isfahan and PD-related human gene expression in the SN of PD patients were also observed (Figure 6). No significant differences were observed in VRFC of these phages across groups, indicating gene expressions of these viruses were comparable between datasets (Supplementary Figure 3). Together, these suggest that phages may disrupt human gene expression in the SN to inhibit dopamine biosynthesis, neural growth factors and anti-viral factors. Symbiotic virobiota may increase the risk of PD.

**Figure 5:**
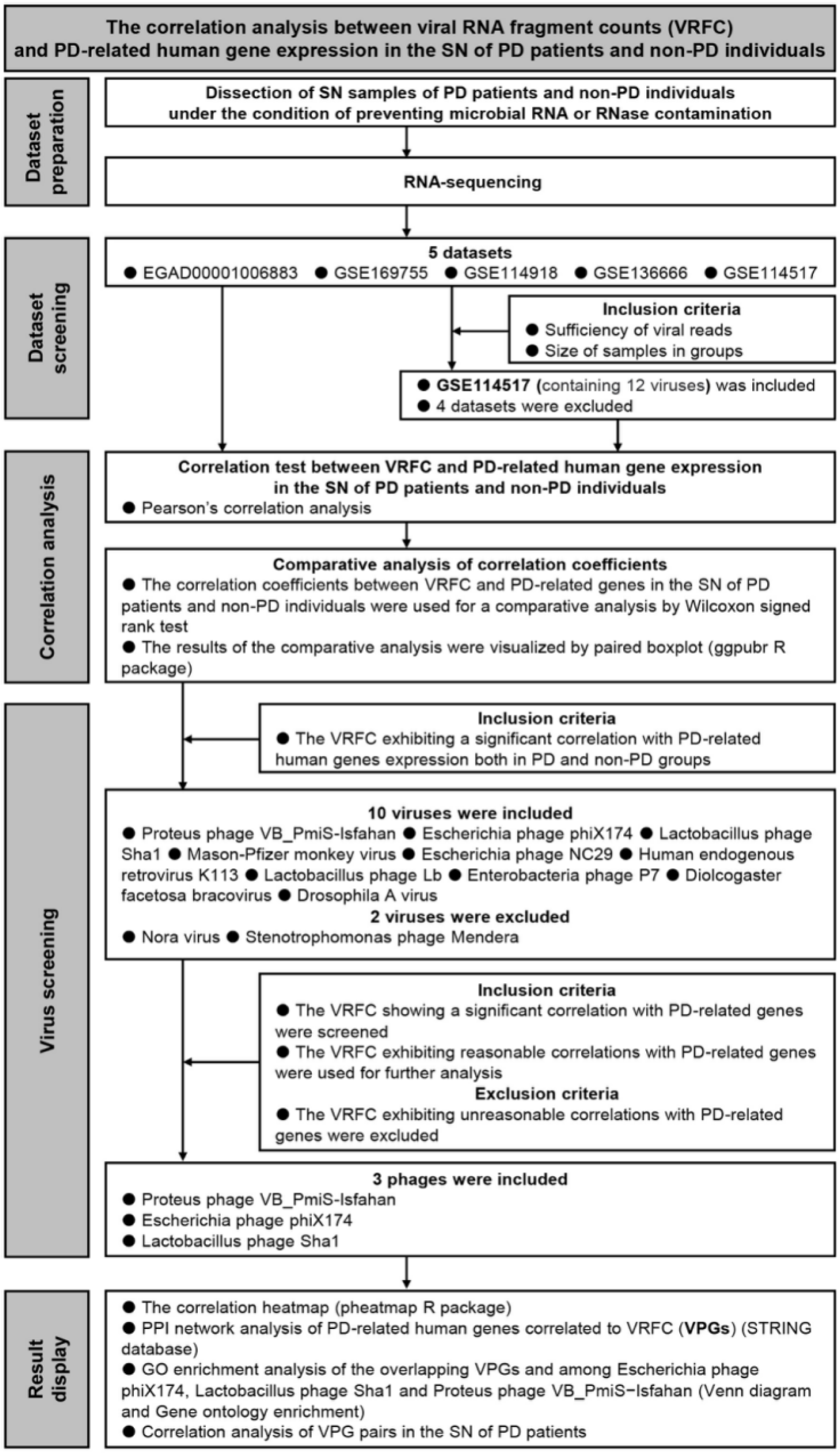
Enrollment of datasets and correlation analysis between VRFC and PD-related human gene expression.

**Figure 6:**
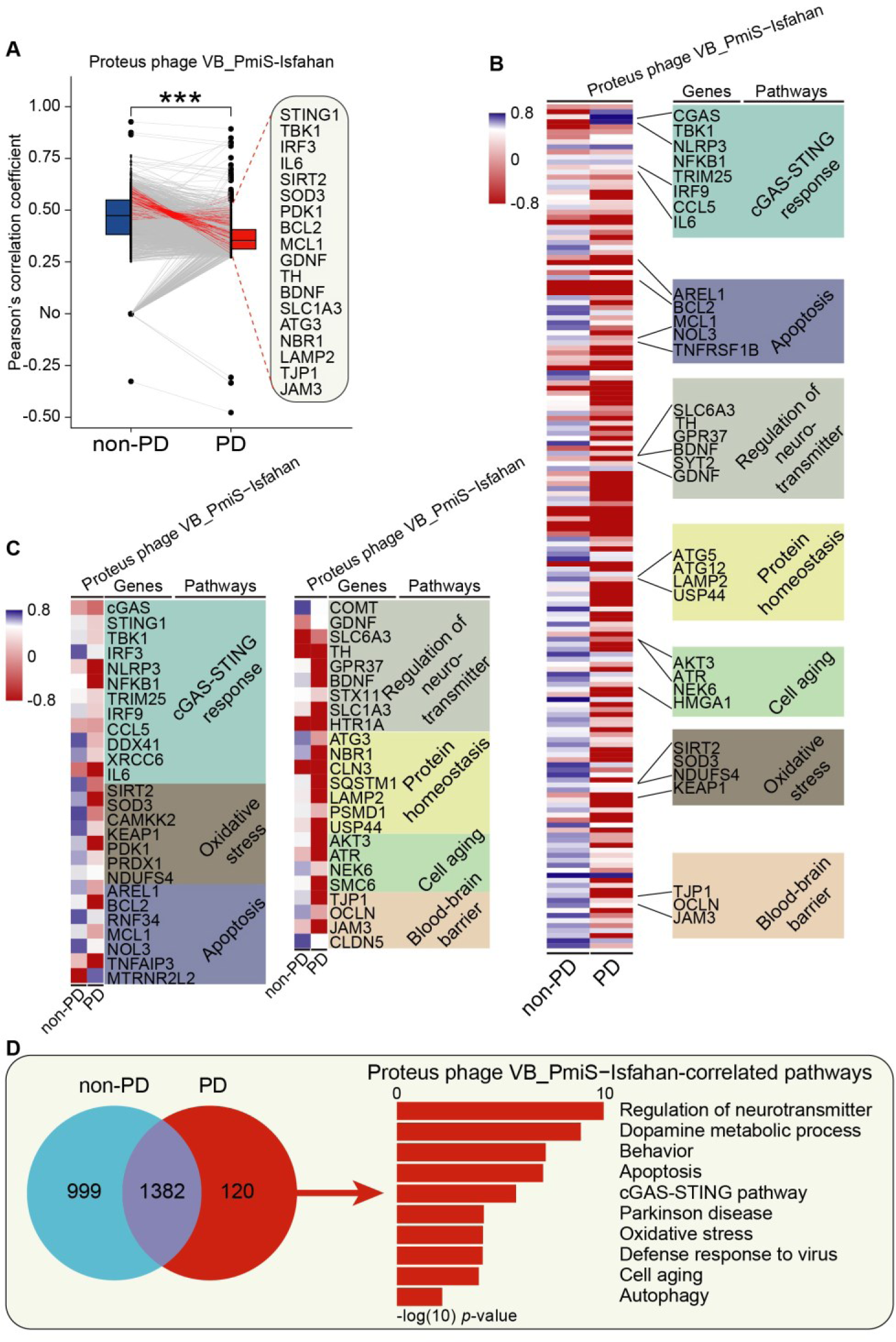
A strong negative correlation between VRFC of Proteus phage VB_PmiS-Isfahan and PD-related human gene expression in the SN of PD patients. (A) Paired boxplots showing changes of the correlations between VRFC of Proteus phage VB_PmiS-Isfahan and PD-related human gene expression in the SN of PD patients and non-PD individuals. Top and bottom edges represent the 1st and 3rd quartiles, respectively, the center line represents the median. The p values were calculated using the Wilcoxon matched-pairs test. ***p value < 0.001. (B) Hierarchical clustered heatmap of correlation profiles between VRFC of Proteus phage VB_PmiS-Isfahan and PD-related human gene expression in the SN of PD patients and non-PD individuals. In the heat map, each column represents a phage, and each row represents a human gene. Red denotes negative correlation, and blue denotes positive correlation. (C) Heatmap of correlation profiles between VRFC of Proteus phage VB_PmiS-Isfahan and expression of PD-related genes in the SN of PD patients and non-PD individuals. (D) The Venn diagram showing the overlapping PD-related genes and pathways between non-PD individuals and PD patients of Proteus phage VB_PmiS-Isfahan.

### DEGs reveal a potential association between symbiotic viruses and PD pathogenesis

Differentially expressed genes (DEGs) reflect molecular signatures of PD. A total of 151 DEGs were identified (Figure 7A and eTable 8). These DEGs were enriched for dopamine biosynthesis, cGAS-STING pathway and response to virus (Figure 7B, C and D). PPI analysis of DEGs revealed similar results with VPGs (Supplementary Figure 4C and D). ROC curve analysis showed that these DEGs can differentiate PD patients from non-PD patients (Supplementary Figure 4E and F), demonstrating that cGAS-STING and antiviral systems may be involved in PD pathogenesis. Venn analysis also highlighted the importance of dopamine biosynthesis and cGAS-STING system in PD etiology (Figure 8). Overall, similar to VPGs, DEGs uncovered a suppressed anti-viral cGAS-STING pathway in the SN of PD patients; validated for the first time that symbiotic virobiota underlies PD pathogenesis; and provide a novel insight into the understanding of PD pathogenesis from the perspective of virus-human symbiosis.

**Figure 7:**
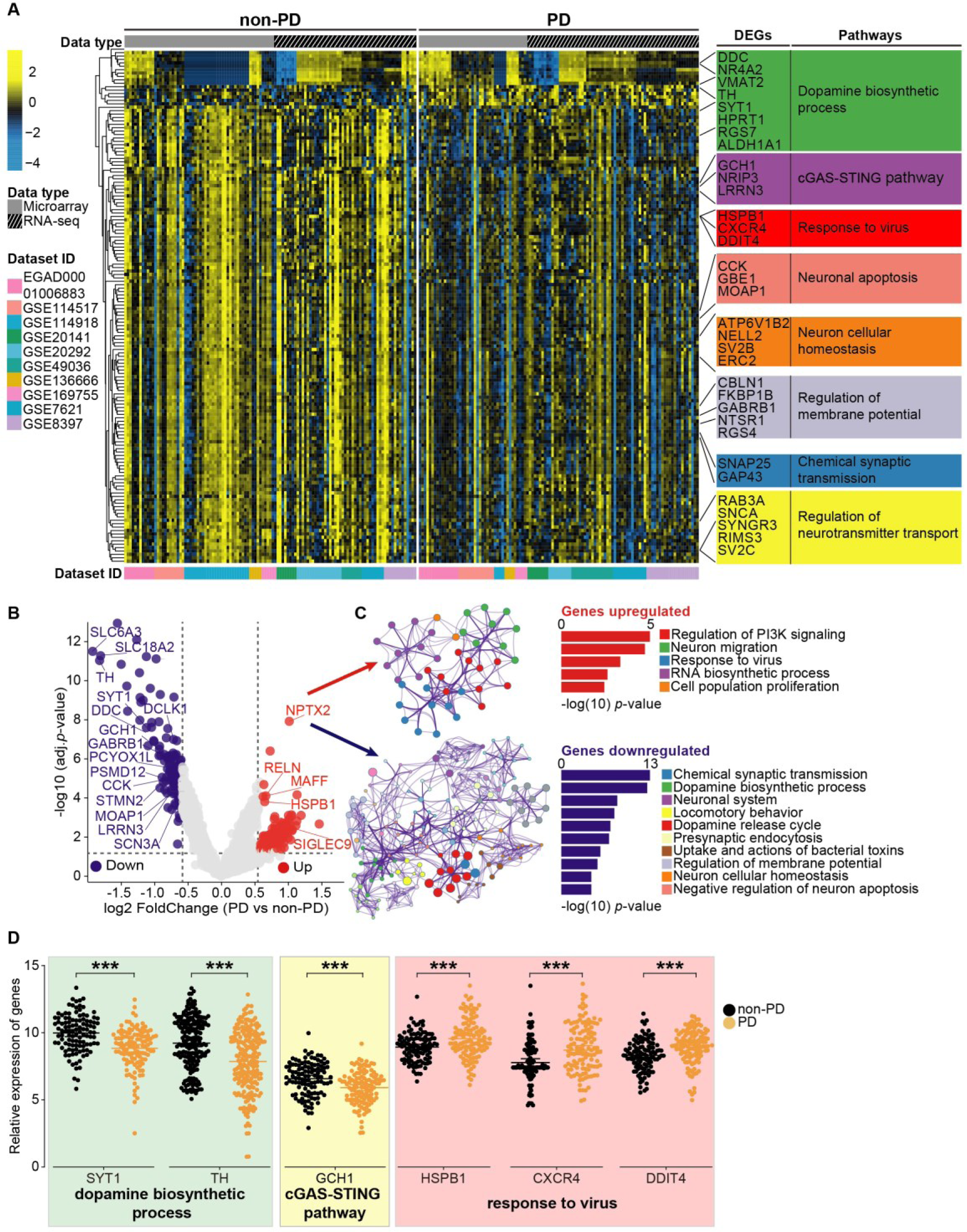
DEGs reveal a potential association between symbiotic phages and PD pathogenesis. (A) Hierarchical clustered heatmap of DEGs in the SN of PD patients and non-PD individuals. (B) Volcano plot displays DEGs in the SN of PD patients compared with non-PD individuals. Upregulated genes are colored in red, downregulated genes are colored in blue, insignificantly altered genes are colored in gray. (C) Gene Ontology and KEGG enrichment analysis of DEGs. (D) The expression level of genes. The total relative expression levels of genes are shown as median and 95% CI. The relative expression level of genes in the SN of PD patients (n = 118) and non-PD individuals (n = 111). Statistical significance was determined by Wilcoxon test, \*\*\**p* value < 0. 001.

**Figure 8:**
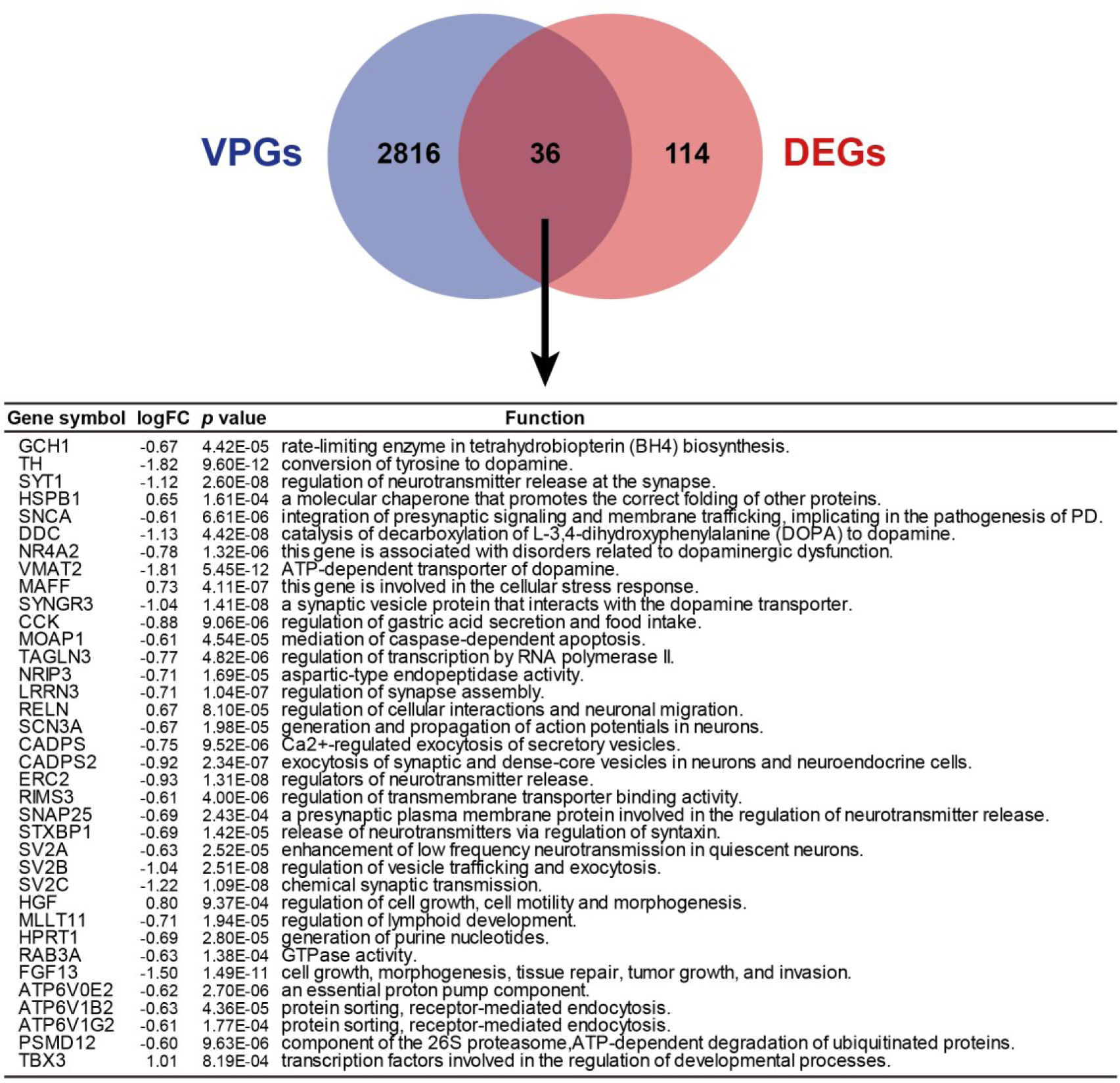
The overlapping genes between VPGs and DEGs in the SN.

**Figure 9:**
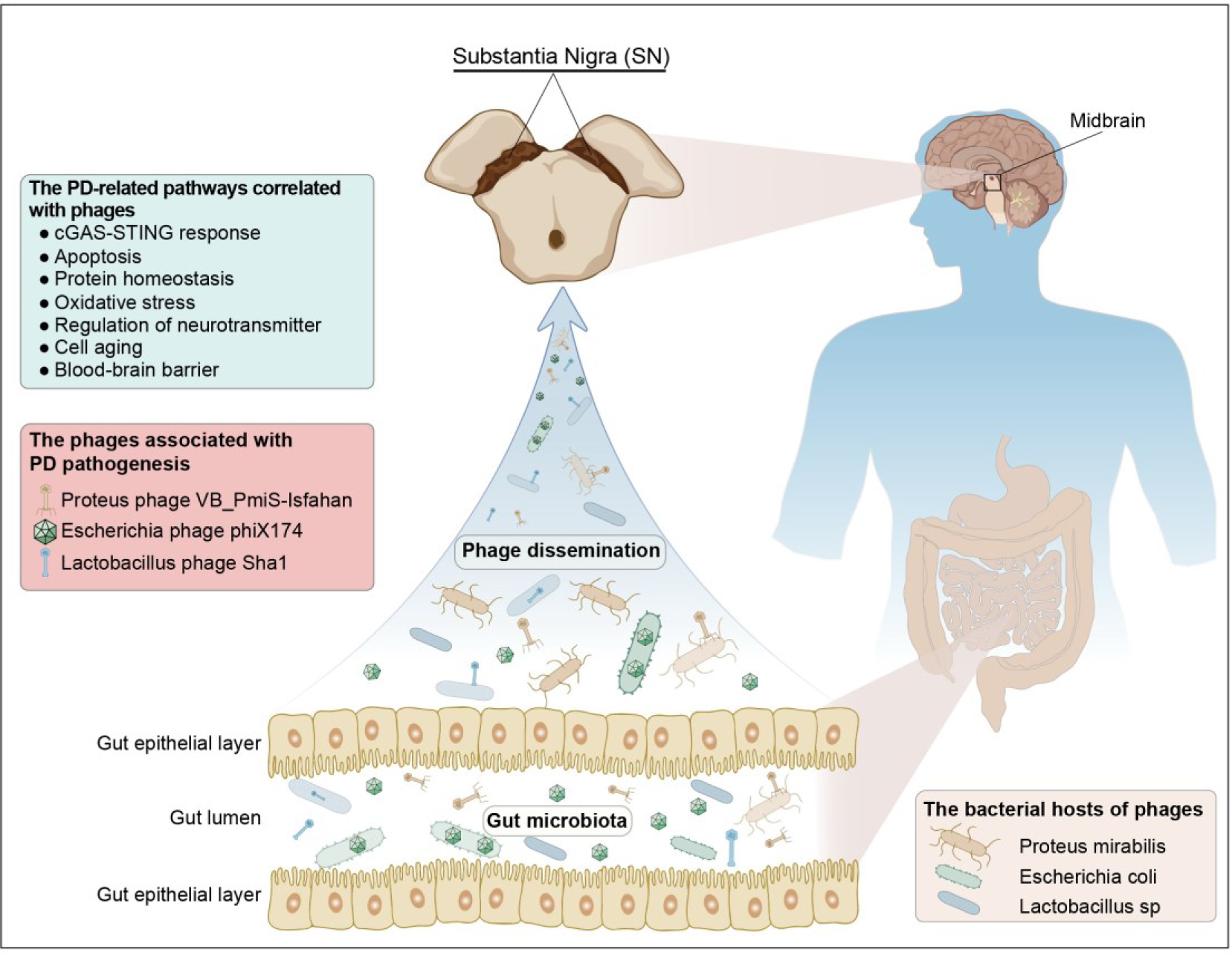
Schematic summary of hypothesized mechanism underlying the virobiota, phages, gut microbiota, and PD pathogenesis. The discovery of symbiotic virobiota in the SN. A symbiotic phages and the dysbiosis of virobiota in the SN may underlie PD pathogenesis. The phages may be one of the bridges between gut microbiota and PD pathogenesis.

## Discussion

For a century, the association between viruses and PD pathogenesis has been debated, but largely ignored or dismissed as controversial^30^. The link between viruses and PD has been defined as one of the medical mysteries, due to clinical studies have shown that there was no history of an overt episode of viral infection in the vast majority of PD patients^28,29^. In 2022, viruses were observed in the amygdala, median temporal gyrus, substantia nigra, intestine and blood of PD patients, showing the positive rates of viruses in PD patients might be higher than that in non-PD individuals^83^. New concepts of ‘Virobiota’^44^ and ‘Virome’^84^ are put forward, revealing a new landscape of virus-human symbiosis, suggesting a potential association between symbiotic virobiota and PD pathogenesis. Nevertheless, limitation of human brain biopsy and difficulty for electron microscope in observing virions have been the major hurdles for deciphering whether or not viruses exist in human substantia nigra^85^.

In this study, using worldwide RNA-sequencing datasets of the SN of PD patients^86–90^, we observed existence of viruses/virobiota in the human SN. We unveiled a dysbiosis in the SN virobiota of PD patients. We found that the phages that host the gut microbiota may be predominant in the SN virobiota of PD patients (Figure 2). These observations suggest that virobiota may exert a life-long influence on neural cells and finally may cause the loss of dopaminergic neurons in the SN. The phages that host the gut microbiota may enter the SN to live in symbiosis with dopaminergic neurons. The phages may be the main bridge between intestinal flora and PD pathogenesis. Together, we show that the virobiota in the SN may underlie PD etiology (Figure 5).

Proteus phage VB_PmiS-Isfahan is a phage that infects Proteus mirabilis^91^, which is one of the dominant human gut bacteria^92^. This phage is present in semen and may be associated with the male fertility^93^. This phage is one of the top abundant viral strains in nasopharyngeal specimens of non-COVID individuals^94^; and it can also be detected in pan-cancer samples^95^. In this study, we found that Proteus phage VB_PmiS-Isfahan was a common virus in the SN of PD patients from the UK, the Netherlands, and Switzerland. The human gene expressions dampened by Proteus phage VB_PmiS-Isfahan were enriched for PD-related pathways including cGAS-STING response, dopamine metabolic process, oxidative stress and apoptosis (Figure 3). Escherichia phage phiX174 is a phage that infects Escherichia coli^96^, which is an indicator of normal intestinal flora of humans. In 1962, Walter Fiers and Robert Sinsheimer identified the phage phiX174 as a single stranded DNA virus^97^. In 1967, Arthur Kornberg used phage phiX174 as the first in vitro model to prove that the synthesized DNA produces all the features of natural virus^98^. As the first DNA-based genome, Frederick Sanger’s group sequenced the genome of phage phiX174 in 1977^99,100^. The phage phiX174 can induce humoral immune response^101^, and thus it has been used to evaluate humoral immune function^101–105^. The phage phiX174 genome can be integrated into human lymphocyte genome^106^. Lactobacillus phage Sha1 is a phage that infects Lactobacillus^107^, and it is increased in the gut virobiota of older adults with mild cognitive impairment^108^. In this study, we observed a strong negative correlation between VRFC of these phages and PD-related human gene expression in the SN of PD patients (eFigure 2). Together, these findings reveal the importance of studying the relationship between phages and etiology of human diseases such as PD.

Dysbiosis of virobiota is involved in the onset and progression of diseases^44,109,110^. Species of phages was decreased, whereas viral community dissimilarity was higher in gut virome of patients with ulcerative colitis than controls^111,112^. There is a difference in gut virome between obese patients with type 2 diabetes mellitus and healthy controls^109^. In this study, we found that virome composition in the SN of PD patients and non-PD individuals was separated into two distinct clusters. The viral community dissimilarity among PD patients was higher than controls. The core species of viruses was higher, whereas unique species of viruses was lowered in the SN of PD patients compared with controls (Figure 2). Overall, these results suggest that dysbiosis of virobiota in the SN may be involved in PD pathogenesis.

Cnaphalocrocis medinalis granulovirus is the most prevalent virus in gut virome of patients with hypertension, whereas this virus is not a contributor to the hypertension^110^. Similarly, in this study, we found that the abundance of Acinetobacter phage AbKT21phiIII and Escherichia phage ESSI2_ev239 was enriched, whereas the abundance of Escherichia phage Lambda_ev017 was diminished in virobiota of the SN of PD patients; and the viral gene expression of these phages was not correlated with PD-related human gene expression. These observations suggest that the abundance of symbiotic viruses in the SN may not be a hallmark of PD pathogenesis, reflecting a complex etiologic connection between symbiotic virobiota and human diseases.

Phages may cause neuroimmune responses of eukaryotic cells. A phage cocktail stimulates IFNγ production in dendritic cells^113^. Staphylococcus phage reduces lipopolysaccharide-induced high levels of IL1β, IL6 in mammary alveolar epithelial cells^114^. Phage lysates up-regulate IL1β and IL6 in peripheral blood mononuclear cells^115^. In this study, we found that viral RNA fragment counts of Proteus phage VB_PmiS-Isfahan, Escherichia phage phiX174 and Lactobacillus phage Sha1, were negatively correlated with the gene expression of proinflammatory cytokines, cGAS-STING pathway, and antiviral immunity. Together, these findings suggest that symbiotic phages in the SN may cause neuroinflammation and disrupt antiviral immune responses, which may contribute to PD pathogenesis.

cGAS–STING system acts as a sensor for cytosol viral DNA upon viral infection and phage invasion^113,116–120^. Activation of cGAS–STING initiates anti-phage immune response to restrict phage activity^121–129^. Virus can inhibit the DNA-sensing function of cGAS-STING system in humans^130^. In this study, we found that both VPGs and DEGs were enriched for cGAS-STING response, and gene expression level of cGAS-STING system was lowered in the SN of PD patients. cGAS-STING system gene expression was negatively correlated with phage gene expression in the SN of PD patients. Together, these findings suggest that cGAS-STING activity may be inhibited by symbiotic phages in the SN of PD patients, revealing a phage-suppressed cGAS-STING function in PD pathogenesis. The relief of phage-inhibited cGAS-STING activity may provide a promising strategy for prevention or treatment of PD.

Epidemiological evidence shows that viral infection may precede the appearance of PD symptoms^131^. In this study, the SN samples were from the PD patients aged between 65 -90 years. Considering the potential symbiosis, we reasoned that the virobiota-neural cell symbiosis in the SN may predate the PD onset.

Amantadine, an anti-Parkinson agent^132^, is also serving as an antiviral medication. Amantadine can inhibit phage assembly^133^. Our previous study suggests that amantadine plays an antiviral role by activating cGAS–STING pathway^134^. These findings raise the possibility that amantadine may eliminate the symbiotic phages to relieve the death of dopaminergic neurons in the SN.

Next-generation sequencing technology can probe viral mRNA fragments of intracellular viruses or viral episomes. RNA-sequencing can detect virobiota in post-mortem SN tissues. In this study, sequencing read length ranged from 50 to 80 bp. The contigs longer than 50 nucleotides showing at least 90% identity to reference viral genome were retained. This strategy guarantees high accuracy and sensitivity of virobiota annotation.

Viral infection may increase BBB permeability. BBB does not constitute a barrier to phages^52–54,135^. Filamentous phage M13 can cross BBB to access the brain after intranasal administration in mice^49,51,136^. Phages can be delivered into the brain through entering peripheral immune cells by the “Trojan horse mechanism”^48,137^. In this study, we found that gene expression of the phages was negatively correlated with BBB-related gene expression (eFigure 2). Together, we suggest that the phages may cross the BBB to symbiose with the SN.

Phage-derived antimicrobials have been broadly applied in clinical treatment, food industry and aquaculture^138–140^. This strategy has been used to treat intestinal, skin, urinary, and respiratory infections^141,142^. Thus, to test whether phage-related therapy may lead to increased risk for PD, further investigations are needed.

Human-gut virome variation is influenced by geographic regions^143^. Our findings revealed that virobiota composition in the human SN was geographically related. Therefore, we suggest that SN samples of PD patients from more countries or regions are needed for assessing association between virobiota and PD pathogenesis.

In summary, this is the first study to discover virobiota in the substantia nigra. A life-long low viral load of symbiotic virobiota in the SN may be a contributor to PD pathogenesis. The phages that host gut microbiota may be implicated in PD etiology. Our observations unlocked the black box between phages and PD, pointing out a complex etiologic connection between symbiotic virobiota and human diseases, providing a novel insight into PD etiology from the perspective of phage-human symbiosis. The further study of virobiota in the brain may shed light on PD pathogenesis and therapy.

## Author Contributions

Yun Zhao and Yuan Zhou did the RNA-seq and microarray data collection. C.X. and Yun Zhao analyzed the data. Yun Zhao, C.X. and R.Z. made the figures. Yun Zhao did the literature search and wrote the paper. Yuan Zhou designed the protocol of bioinformatic analysis. C.X. performed the bioinformatic analysis. B.W., D.L., J.L., S.W., and Y.H. participated in the study. R.Z. conceived the study, wrote and edited the paper. All authors contributed to the study design, reviewed, edited and approved the manuscript for submission. All authors accept full responsibility for the content of this paper. R.Z. is the lead contact.

## Funding

This work was supported by grants from the National Natural Science Foundation of China (No. 82170864. No. 81471064. No. 81670779 and No. 81870590 to R.Z), the National Key Research and Development Program of China (2017YFC1700402 to R.Z.), the Beijing Municipal Natural Science Foundation (No. 7162097 and No. H2018206641 to R.Z), the Peking University Research Foundation (No. BMU20140366 to R.Z), Scientific Project of Beijing Life Science Academy (No. 2023300CB0100 to R.Z).

## Acknowledgments

We thank Paul Lingor and Lucas Caldi Gomes (Department of Neurology, Rechts der Isar Hospital, Technical University of Munich, München, Germany) for kind permission and assistance in acquiring the EGA data. We thank Chunfu Zheng (Department of Microbiology, Immunology & Infection Diseases, University of Calgary, Calgary, Alberta, Canada) for helpful comments and discussions of this manuscript. We thank Hermona Soreq (Department of Biological Chemistry, The Alexander Silberman Institute of Life Sciences, The Hebrew University of Jerusalem, Jerusalem, Israel), Eva Hedlund (Department of Biochemistry and Biophysics, Stockholm University, Stockholm, Sweden) for discussions about the read length of RNA-seq. We thank Yingfei Ma (CAS Key Laboratory of Quantitative Engineering Biology, Shenzhen Institute of Synthetic Biology, Shenzhen Institutes of Advanced Technology, Chinese Academy of Sciences, Shenzhen, China) for helpful comments about virus classification. We thank Weiguang Zhang, Lihua Qin, Ke Wang (Department of Anatomy, Histology and Embryology, Peking University, China), Fuping You (Department of Immunology, Peking University, China) and Yangyi Fan (Department of Neurology, Peking University People’s Hospital) for technical support. We thank Hui Yang (Department of Neurosurgery, Huashan Hospital, Institute for Translational Brain Research, Shanghai Key laboratory of Brain Function Restoration and Neural Regeneration, Shanghai Clinical Medical Center of Neurosurgery, MOE Frontiers Center for Brain Science, Shanghai Medical College, Fudan University, Shanghai, China) for providing the gene list of cGAS-STING system.

## Data Availability Statement

The raw RNA-seq data in this study were retrieved from NCBI-GEO (http://www.ncbi.nlm.nih.gov/geo/) or generously shared by Prof. Paul Lingor and Dr. Lucas Caldi Gomes on European Genome-phenome Archive (EGA) platform (https://ega-archive.org/). The corresponding accession numbers: EGAD00001006883, GSE169755, GSE114918, GSE136666 and GSE114517. The raw microarray in this study were retrieved from NCBI-GEO. The corresponding accession numbers: GSE20141, GSE49036, GSE7621, GSE8397 and GSE20292. Details of the datasets are provided in Table S1. The information of PD patients and non-PD individuals in this study are provided in Table S2.

## Declaration of Competing Interest

The authors declare that they have no competing interests.

## Notes

### Competing Interest Statement

The authors have declared no competing interest.

