## Supplemental figures for "The discovery of phages in the Substantia Nigra and its implication for Parkinson’s Disease"

### Legends of supplemental figures

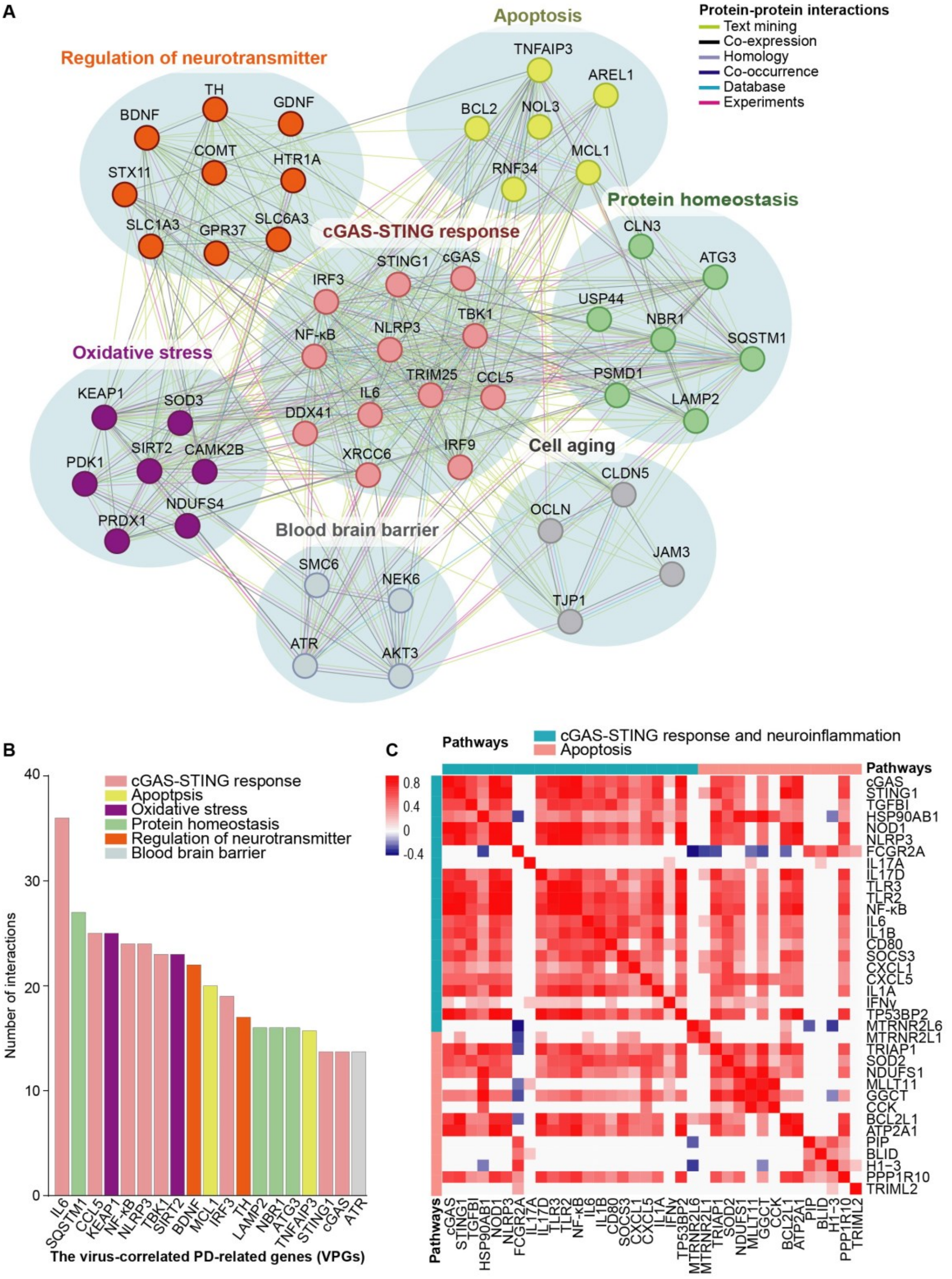


**Figure S1: PPI network analysis and correlation profiles between cGAS-STING response, neuroinflammation and apoptosis in the SN of PD patients**

(A) PPI network analysis of VPGs. The genes with more than two interactions are shown in the network. (B) Hub node genes determined by interaction numbers. A total of 15 genes with more than 10 interactions are shown in the bar plot. Abbreviation: Abbreviation: IL6, interleukin 6; SQSTM1, sequestosome 1; CCL5, C-C motif chemokine ligand 5; KEAP1, kelch like ECH associated protein 1; NF-κB, nuclear factor kappa B subunit 1; NLRP3, NLR family pyrin domain containing 3; TBK1, TANK binding kinase 1; SIRT2, sirtuin 2; BDNF, brain derived neurotrophic factor; MCL1, myeloid cell leukemia-1; IRF3, interferon regulatory factor 3; TH, tyrosine hydroxylase; LAMP2, lysosomal associated membrane protein 2; NBR1, neighbor of BRCA1 gene 1; ATG3, autophagy related 3; TNFAIP3, TNF alpha induced protein 3; STING1, stimulator of interferon response cGAMP interactor 1; cGAS, cyclic GMP-AMP synthase; ATR, ATR serine/threonine kinase. (C) Heatmap and clustering of VPGs based on their gene-gene pair correlations. Rows and columns represent human genes. (red: positive correlation; blue: negative correlation).

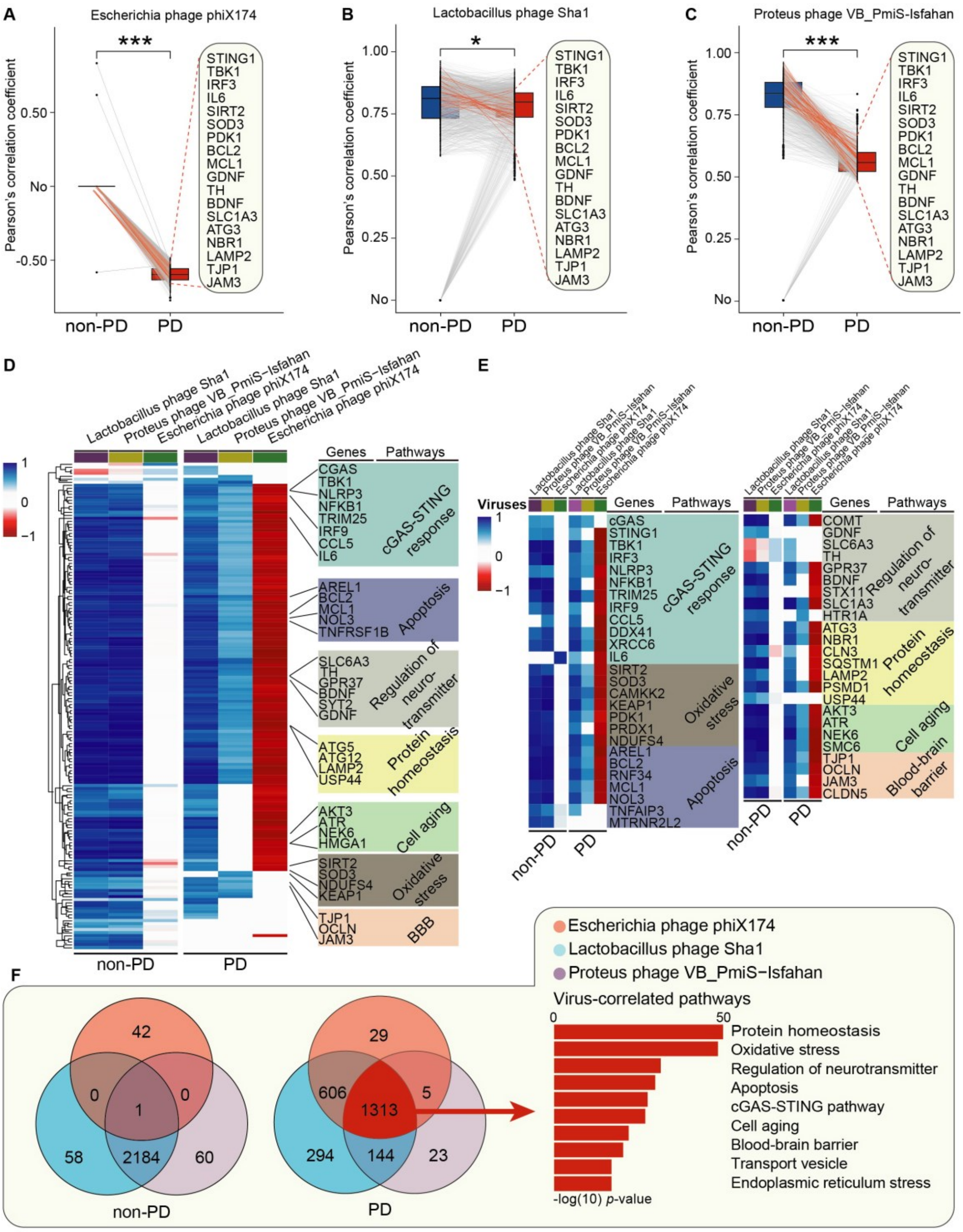


**Figure S2: A strong negative correlation between VRFC of the phage phix174 and PD-related human gene expression in the SN of PD patients**

(A, B, C) Paired boxplots showing changes of the correlations between viral RNA fragment counts (VRFC) of phages and PD-related gene expression in the SN of PD patients and non-PD individuals. Top and bottom edges represent the 1st and 3rd quartiles, respectively, the center line represents the median. The p values were calculated using the Wilcoxon matched-pairs test. *p value $<$ 0.05 and ***p value $<$ 0.001. (D) Hierarchical clustered heatmap of correlation profiles between VRFC of phages and PD-related human gene expression in the SN of PD patients and non-PD individuals. In the heat map, each column represents a phage, and each row represents a human gene. Red denotes negative correlation, and blue denotes positive correlation. (E) Heatmap of correlation profiles between VRFC of phages and expression of PD-related genes in the SN of PD patients and non-PD individuals. (F) The Venn diagram showing the overlapping PD-related genes and pathways among Escherichia phage phiX174, Lactobacillus phage Sha1 and Proteus phage VB_PmiS-Isfahan.

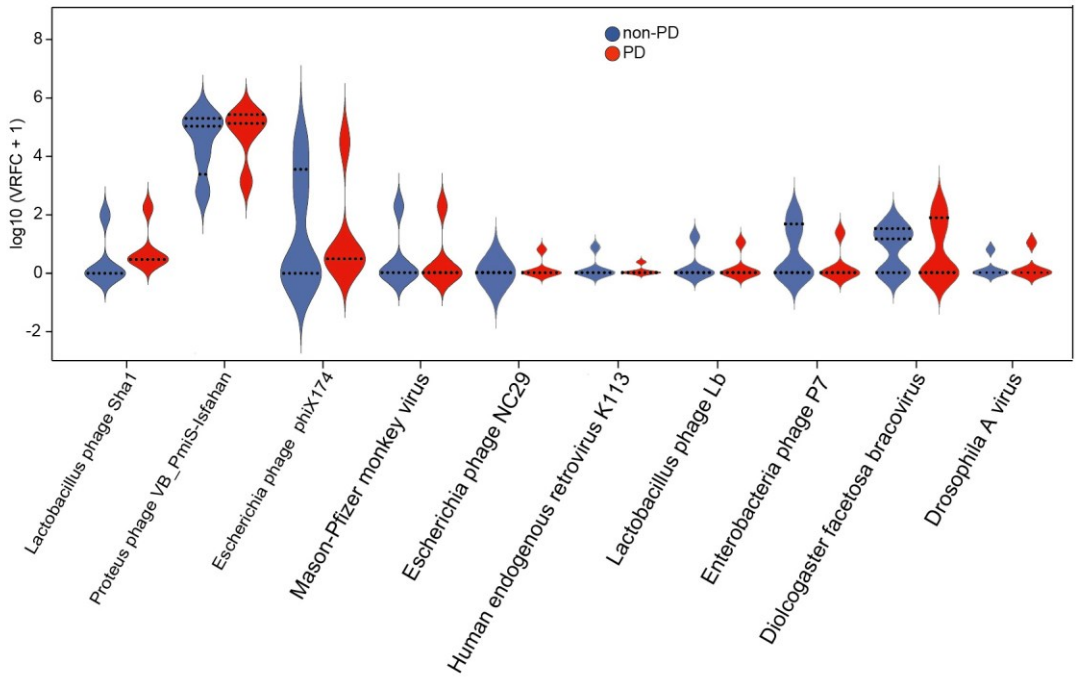


**Figure S3: Comparison of virobiota abundance in the SN of PD patients and non-PD individuals**

Violin plot showing the $\log_{10}$(VRFC + 1) of viruses in the SN of PD patients and non-PD individuals.

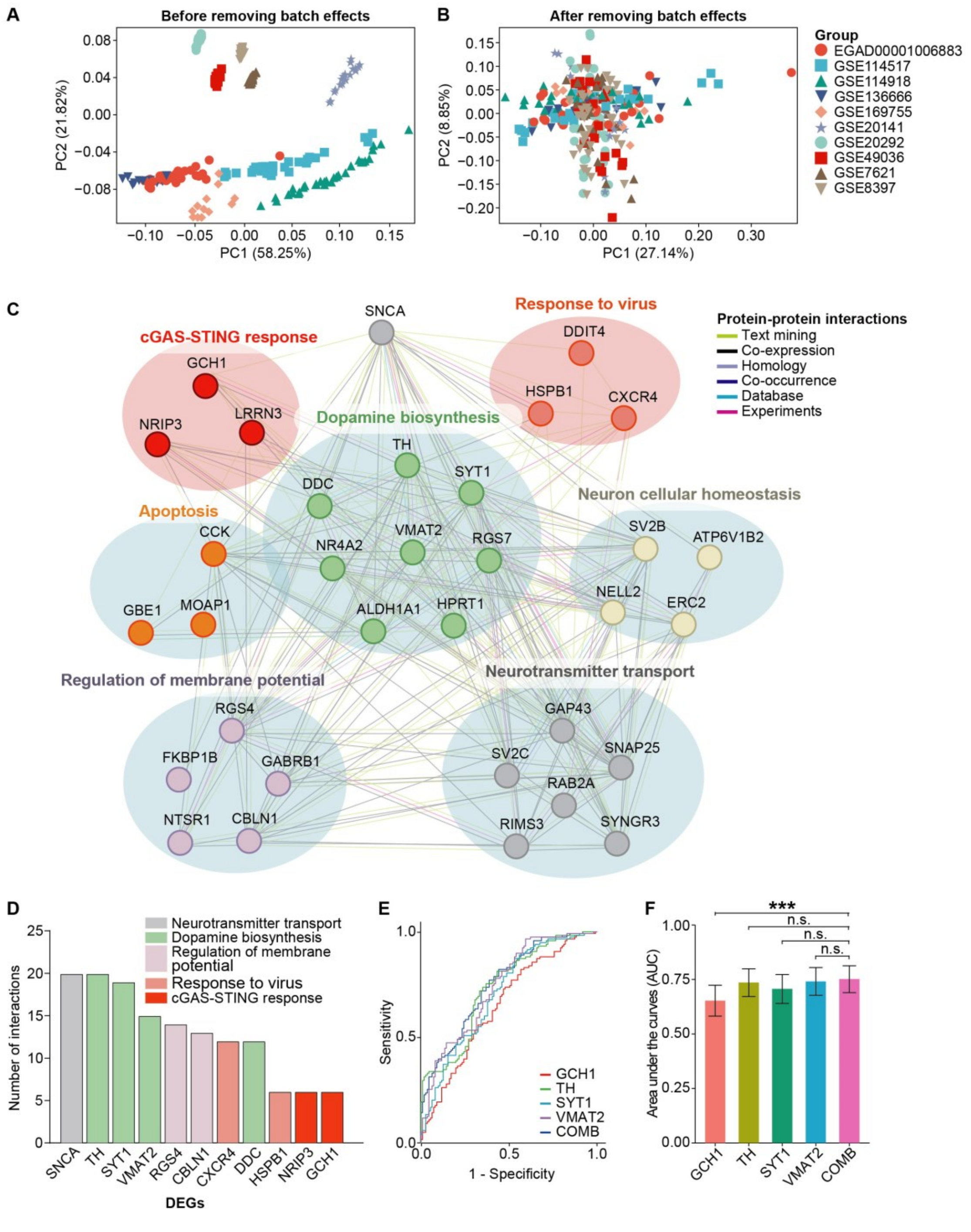


**Figure S4: PPI and ROC curve analysis of DEGs**

Principle component analysis (PCA) plot before removing batch effects (A) and after removing batch effects (B) for the RNA-seq datasets and microarray datasets. (C) PPI network analysis of DEGs. (D) Hub node genes determined by interaction numbers. A total of 24 genes with more than 5 interactions are shown in the bar plot. (E) ROC curve analysis of each and combination (COMB) of hub node genes. The curve of combination of four genes lies in the highest position than that of GCH1, TH, SYT1, and VMAT2 alone. (F) AUCs of each and combination of hub node genes. n.s. not significant; ***$p$ value $<$ 0.001.
